# Prebreeding populations and the importance of life history for conserving the world’s imperiled seabirds

**DOI:** 10.1101/2025.02.19.639124

**Authors:** Liam U. Taylor, Eleanor Gnam

## Abstract

Seabird conservation often focuses on nestlings and breeding adults. Yet imperiled seabird populations also contain prebreeders, including juveniles and subadults, that wait several years before breeding at colonies. We use previously-published data on reproductive and survival rates for 84 species to quantify the conservation relevance of prebreeding seabirds. We find, first, that prebreeders average about half of seabird populations (median 47.4%, range 11.2%–66.7%). Second, while seabird population growth is much more sensitive to adult survival than prebreeder survival, human-driven changes may shift the importance of prebreeders for future population stability. Third, lowering the breeding age is a powerful, but underexplored, route to increasing population growth. Managing prebreeders could thus play a key role in protecting seabirds. This task may require answering fundamental questions about the behavior of young birds. Broadly, we suggest that life history characteristics (e.g., breeding age) actively shape both obstacles to, and opportunities for, successful conservation.

## Introduction

Many seabird populations are in precipitous decline (Croxall *et al*. 2012). From gulls to albatrosses, hundreds of species of coastal and pelagic birds face dozens of threats, including overfishing that depletes food supplies, invasive predators that jeopardize survival, and warming waters that disrupt ecosystems (Dias *et al*. 2019). New threats, such as outbreaks of deadly avian influenza, continue to emerge (Klaassen and Wille 2023).

In this grim context, many conservationists labor to protect seabird populations. Examples of important management efforts include eradicating predators near breeding grounds (Bird *et al*. 2024), installing artificial boxes to protect nests (Libois *et al*. 2012), and using social attraction to build populations (Spatz *et al*. 2023). Because these management efforts take place at breeding colonies, they mainly protect the nests and breeding adults within those colonies.

Yet breeding colonies host only a portion of seabird populations (Ainley *et al*. 2024). Nearly all seabird populations include young birds that have not yet begun reproducing at colonies (“prebreeders”). Seabirds typically delay their first breeding attempt for several years, sometimes up to a decade (Lack 1968). Therefore, prebreeding populations include both juveniles in their first year of life, which is typically spent at sea, and subadults that spend subsequent years traveling, foraging, and visiting colonies before settling down to breed.

Decades ago, biologists began asking questions about the behavior of young seabirds (Wynne-Edwards 1962; Ashmole 1963). Recently, miniaturized tracking devices have renewed interest in young marine animals, including seabirds, during their “lost years” at sea (e.g., Collet *et al*. 2020; Phillips *et al*. 2025; Table S1). However, most prebreeding populations remain unmonitored, unprotected, and understudied.

It is conventional wisdom that seabirds have “slow” life histories, with small clutches and high survival rates (Lack 1968). Consequently, the survival of breeding adults is expected to be more important for population stability than is the survival of juveniles or subadults (Sæther and Bakke 2000). In practice, however, unique management actions that reduce the mortality of young seabirds can supplement population growth (Finkelstein *et al*. 2010). Prebreeders may also help stabilize populations when they return to colonies and begin breeding at atypically young ages, as has been observed in low-density years following disease outbreaks (Sceviour *et al*. 2024) and culls (Coulson *et al*. 1982). These observations suggest that the survival and breeding age of young seabirds could play key roles in overall population dynamics.

Here, we use simple demographic models to quantify multiple ways in which prebreeding populations could influence conservation outcomes. Specifically, we analyzed previously-published breeding and survival rates across 84 seabird species to answer three questions: (1) What proportion of total seabird populations are prebreeders? (2) How sensitive is future population growth to changes in prebreeder survival? (3) How sensitive is future population growth to changes in age at first breeding?

## Methods

### Data collection

We compiled demographic parameters from published literature on seabird populations. Our analyses required the following five parameters for each species: age at first breeding (in years), annual fecundity, juvenile survival rate, annual subadult survival rate, and annual adult survival rate. Fecundity was calculated as breeding propensity (i.e., the probability of nesting each year) times productivity (i.e., average number of female fledglings per nest, assuming equal sex ratios).

We defined juveniles as birds in their first year, from fledging to 12 months old, and subadults as birds between 12 months old and age at first breeding. Accordingly, juvenile survival rate covered the period from fledging to 12 months old. Subadult survival rate covered the period from 12 months to age at first breeding.

Our initial search for demographic data included 363 seabird species in six taxonomic orders (Table S2). First, we referenced each species account in Cornell’s Birds of the World (Billerman *et al*. 2022). We then queried Google Scholar (e.g., “<SCIENTIFIC NAME> <COMMON NAME> mark recapture survival”) to find additional studies with demographic data from marked populations. Whenever possible, we used female demographic rates from a single long-term study population.

The format and precision of demographic parameters varied among species and sources. For example, subadult survival rate was variously provided as a separate parameter, derived from overall survival rate from fledging to breeding age (n = 18), or assumed to be identical to adult survival (n = 14). Breeding propensity was often unavailable when all other parameters were available. In such cases (n = 30 species), we estimated breeding propensity as the mean breeding propensity of species in the same family in our dataset (Fig. S1A). The final dataset totaled 84 species, spanning six orders and 12 families (Table S2). Details, justifications, and citations for each demographic parameter are provided in the supplemental data file.

### Demographic analysis

For each species, we constructed a simplified life table with three age classes: juveniles, subadults, and breeding adults. We formatted this information as a Leslie matrix—a compact representation of population-level breeding and survival rates—which allowed us to project population growth through time (Caswell 2001 pp. 20–33). We conducted all analyses in R v4.4.1 (R Core Team 2024). See Panel S1 for technical details on Leslie matrices and simplifying assumptions.

First, we used stable age distributions to estimate the proportion of each seabird population that consists of prebreeders. The stable age distribution gives the expected ratio of individuals in different age classes (i.e., juvenile, subadult, adult). This distribution is calculated as the right eigenvector of a Leslie matrix (Caswell 2001 pp. 86–7). We summed the proportions of juveniles and subadults in each stable age distribution to obtain overall proportions for prebreeders.

Second, we estimated the relative importance of prebreeder survival on future population growth rate using elasticity values. We calculated population growth rate as the dominant real eigenvalue of each Leslie matrix (Caswell 2001 pp. 85–6). Each parameter in the matrix (e.g., fecundity, adult survival rate) has a sensitivity value, which is the effect of a small unit change on population growth rate. Normalizing sensitivity values—proportional to the magnitude of each parameter—gives an elasticity for each parameter. A higher elasticity indicates a greater relative contribution to population growth (Caswell 2001 pp. 206–31).

Because our study used highly simplified demographic data, there were only two elasticity values for each species: one for adult survival, and one covering fecundity, juvenile survival, and subadult survival (Caswell 2001 pp. 231–2). We compared these two values to quantify the relative importance of prebreeder (i.e., juvenile and subadult) and adult survival rates. These elasticities were calculated from the original demographic values in our dataset, but current and future changes to seabird populations may shift these values. For example, management efforts at colonies may improve fecundity (e.g., protecting eggs and chicks with artificial nest boxes; Libois et al. 2012) while threats at sea may reduce adult survival (e.g., fishing vessel bycatch; Dias *et al*. 2019). To illustrate the potential consequences of these anthropogenic changes, we recalculated elasticities across a range (10%–190%) of altered fecundity and adult survival values.

Third, we estimated the sensitivity of future population growth to potential changes in age at first breeding. Breeding age is not a parameter in the Leslie matrix itself but rather influences how the matrix is arranged (Panel S1). Thus, we quantified sensitivity as the hypothetical change in population growth rate given a one-year decrease in breeding age. This simulation does not include potential tradeoffs from younger breeding (Panel S1). Following Stearns (1992), we contextualized this metric by asking: what increase in fecundity would match the population benefit of a younger breeding age?

## Results

### What proportion of seabird populations are prebreeders?

We estimated the proportion of birds in different age classes—juvenile, subadult, and adult—using the stable age distribution calculated from the life table for each species. Across all 84 species, juveniles made up a median of 16.6% (range 4.1%–44.1%) of overall populations, subadults made up 26.6% (7.1%–50.2%), and adults made up 52.6% (33.3%–88.8%). Across species, the median proportion of prebreeders (juveniles plus subadults) was 47.4% (11.2%– 66.7%; Fig. 1).

**Figure 1.**
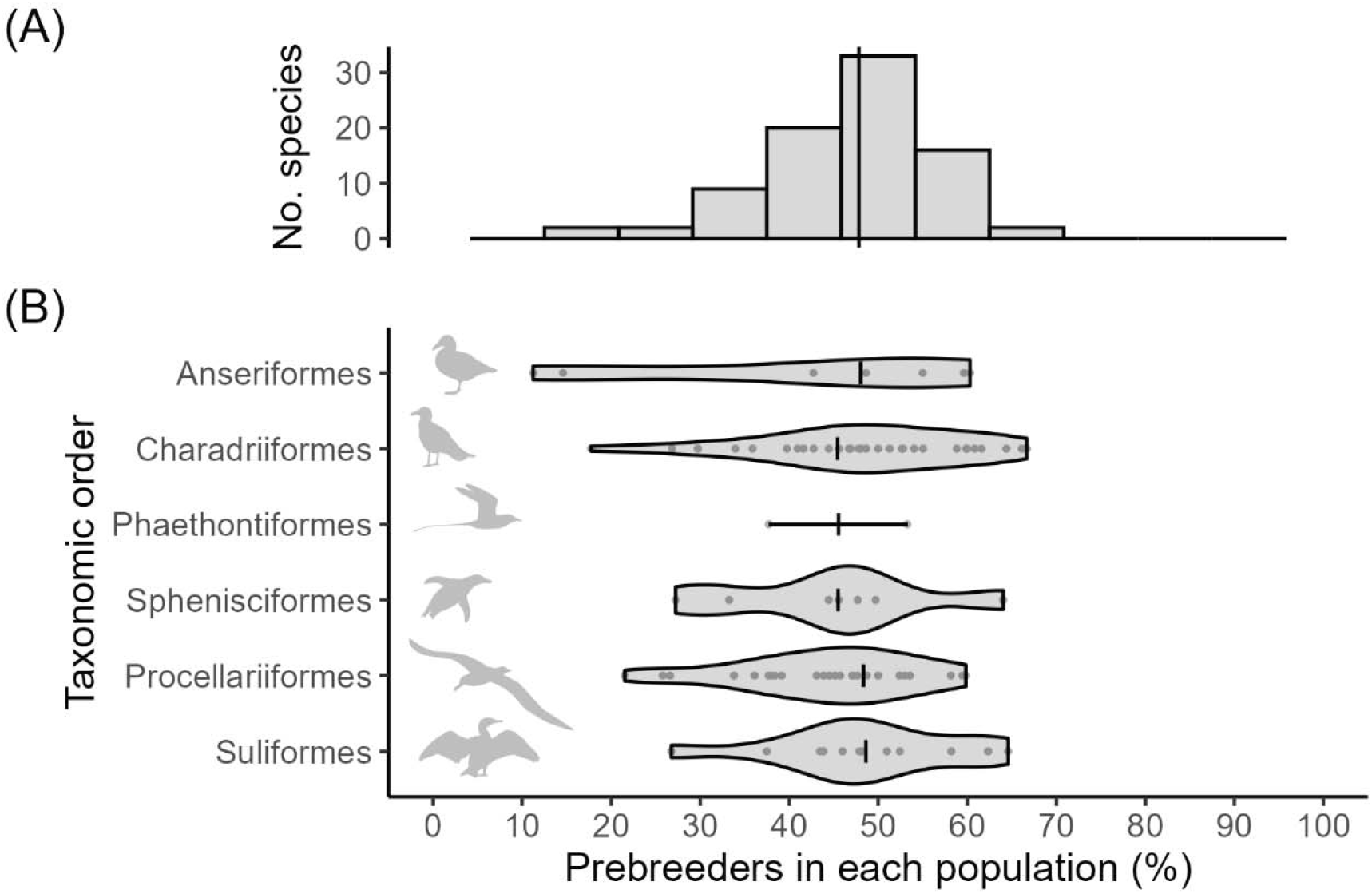
Prebreeders make up large proportions of overall seabird populations. (A) Proportion of prebreeders, including both juveniles and subadults, estimated from the stable age distributions of 84 species. (B) Organized by taxonomic order. Vertical lines show medians. Silhouettes from https://www.phylopic.org/.

### How sensitive is population growth to prebreeder survival?

We calculated elasticities (i.e., relative sensitivities) of different demographic parameters, which quantify the relative impact that changing those parameters would have on future population growth rate. Seabirds had much higher elasticity values for adult survival rate (median 0.61, range 0.40–0.89; Fig. 2A) compared to the alternate elasticity value, which covered all of fecundity, juvenile survival rate, and subadult survival rate (median 0.07, range 0.02–0.19; Fig. 2A).

**Figure 2.**
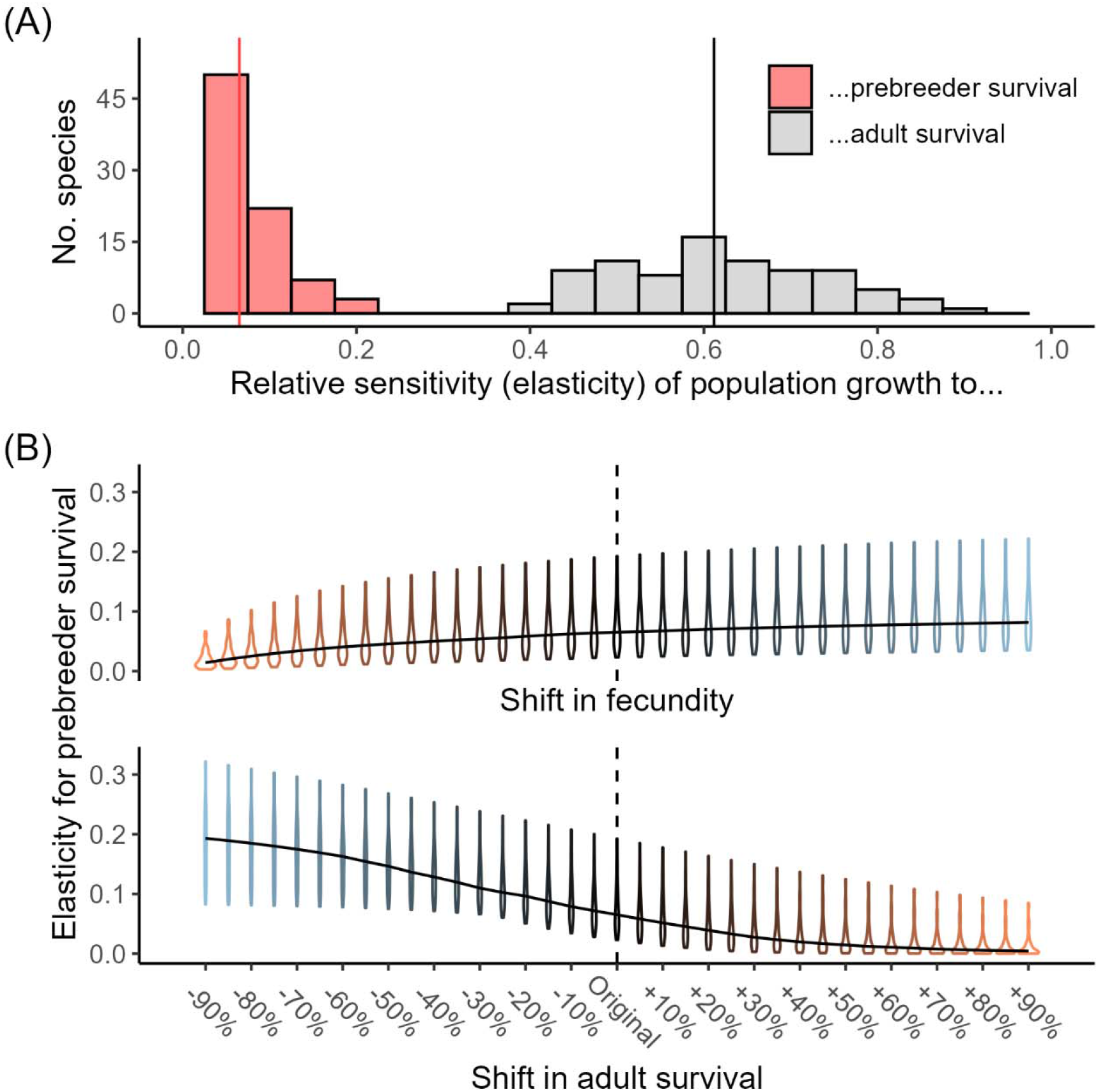
Seabird population growth is far more sensitive to changes in adult survival than prebreeder survival, but future demographic changes can shift these sensitivities. (A) Relative sensitivity (i.e., elasticity) of population growth rate to changes in adult survival rate (gray) or prebreeder survival rates (red) across 84 seabird species. A single elasticity value applies to both juvenile and subadult survival rates (along with fecundity). Vertical lines show medians. (B) Shifting elasticities across a range of alternate values for fecundity or adult survival. Each violin spans 84 species, showing the distribution of relative sensitivity (i.e., elasticity) values for prebreeder survival. Solid black curves show medians.

To illustrate how potential anthropogenic changes may affect seabirds, we recalculated elasticities across a range of altered fecundity and survival values (Fig. 2B). Unsurprisingly, elasticity for prebreeder survival increased slowly whenever fecundity grew, and increased more sharply whenever adult survival declined (Fig. 2B). Absolute effects were small: median elasticity for prebreeder survival never exceeded 0.193 (Fig. 2B), remaining smaller than the original elasticities for adult survival (Fig. 2A). However, relative effects were potentially large within the observed range of ongoing, anthropogenic impacts. For example, a recent study of Arctic Skuas (*Stercorarius parasiticus*) shows a 17% decline in adult survival (from 0.93 to 0.77; Snell *et al*. 2025). Modeling this decline across our dataset raised median elasticity for prebreeder survival from 0.066 to 0.091 (+38%).

### How sensitive is population growth to age at first breeding?

Median age at first breeding in our dataset was four years (Fig. S1B), ranging from two years (as in some sea ducks, such as Spectacled Eider *Somateria fischeri*) to eleven years (as in some large tubenoses, such as Snowy Albatross *Diomedea exulans*). Simulating a one-year decrease in breeding age resulted in a small, but universal, benefit to population growth (Fig. 3A). Median increase in population growth rate was 0.014 (range 0.0005–0.119). For six species, a one-year decrease in breeding age was sufficient to shift from a slowly shrinking population to a slowly growing population (starting growth rates 0.961–0.995, ending growth rates 1.010– 1.060, median increase 0.035; Fig. 3A). The largest benefits from younger breeding—nearing or exceeding a 10% boost in population growth—occurred for species with lower initial breeding ages (Fig. S2). The population benefit from a one-year decrease in breeding age was equivalent to a median 19.5% (1.5%–85.0%) increase in fecundity (Fig. 3B).

**Figure 3.**
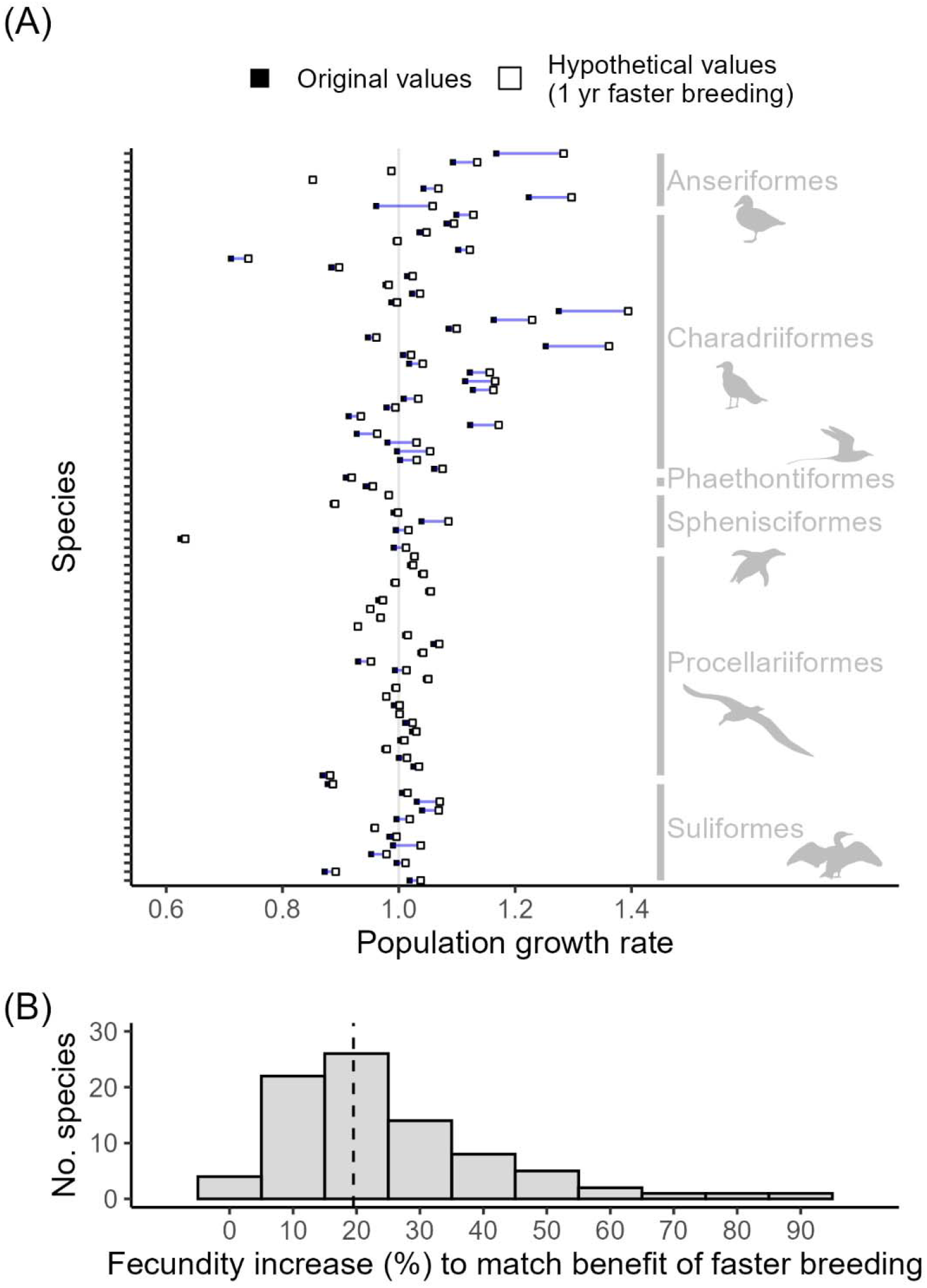
For seabirds with delayed reproduction, lowering the breeding age could increase population growth rate. (A) Hypothetical benefit to population growth rate from a younger breeding age. Closed points show initial population growth rate for each of 84 seabird species. Open points show population growth rate given a one-year decrease in age at first breeding. Blue lines indicate the increase in population growth rate (longer blue lines = bigger benefit). Silhouettes from https://www.phylopic.org/. (B) Percent increase in fecundity needed to match the population benefit of a one-year decrease in breeding age.

## Conclusions

We analyzed 84 species to highlight three ways in which young prebreeders could play crucial roles in the ongoing conservation of seabirds. Our analyses relied on simple demographic assumptions (Panel S1) and did not examine individual species. Instead, we aimed for clear, broad, and explicit conclusions likely to apply across seabirds.

First, prebreeders can make up nearly half of seabird populations (Fig. 1). Many direct conservation actions for seabirds take place at breeding colonies, where managers can, for example, eradicate invasive predators to safeguard nests and breeding adults (Bird *et al*. 2024). Prebreeders are not associated with nests, meaning they may not be represented in colony surveys or receive immediate benefits from colony management (Ainley *et al*. 2024). Similarly, strategies that safeguard birds far offshore—such as the establishment of marine protected areas—are informed by movement patterns of individually tracked birds, which are often breeding adults (Thaxter *et al*. 2012). But recent tracking studies show that prebreeders often travel and forage differently than breeding adults (Table S1). In sum, if prebreeders make up half of seabird populations, then current efforts may offer relatively limited protection for half of seabird populations.

Second, we demonstrated that future anthropogenic changes could make seabird populations increasingly sensitive to prebreeder survival (Fig. 2). Because most seabirds have relatively high survival rates, low fecundity, and delayed reproduction, their population dynamics are most sensitive to mortality amongst breeding adults (Sæther and Bakke 2000). Our analyses show that, on average, seabird population growth is more than six times as sensitive to adult survival than to fecundity, juvenile survival, or subadult survival (Fig. 2A).

However, ongoing changes to seabird population dynamics may increase the importance of prebreeders (Fig. 2B). If, for example, invasive predators start eating adults at a breeding colony (Bird *et al*. 2024), then the stability of that population becomes more reliant on prebreeder survival (Fig. 2B). Management for increased prebreeder survival would benefit populations less than equivalent increases in adult survival (Fig. 2). In practice, however, there may be no such tradeoff. Efforts to protect juveniles and subadults can offer unique opportunities to grow a population (e.g., Finkelstein *et al*. 2010), which may become increasingly valuable as the importance of prebreeder survival increases.

Third, we highlighted how a younger breeding age could lead to greater population growth in seabirds (Fig. 3). A one-year decrease in breeding age increased population growth rate by an average of 0.014, with extremely small gains for some species (e.g., <0.001 for the globally endangered Sooty Albatross *Phoebetria fusca*) but large gains for others (e.g., 0.057 for the US endangered Roseate Tern *Sterna dougallii*; Fig. 3A). On average, younger breeding offered the same benefit as a 19.5% increase in fecundity (Fig. 3B). A well-established idea from evolutionary theory is that, all else being equal, younger breeding leads to increased population growth (Stearns 1992). This idea has practical consequences for seabirds.

The problem is that we do not actually know why seabirds delay reproduction in the first place. Hypotheses include the need for young birds to develop foraging skills (Ashmole 1963) or social behaviors, territories, and mates (Taylor 2024). One important clue is that seabirds may breed at younger ages in lower-density colonies (Sceviour *et al*. 2024, Coulson *et al*. 1982). This observation suggests that management for large, dense colonies may protect breeding adults (Coulson 2001) only at the cost of suppressing the breeding rate of young birds.

In the meantime, important efforts already rely—sometimes implicitly—on manipulating the behavior of young seabirds. It is now commonplace to establish low-density breeding colonies through social attraction via audiovisual decoys (Spatz *et al*. 2023). To the extent that these efforts draw new recruits, rather than just shuffling regional populations around, they must attract birds that are not already breeding elsewhere (Ainley *et al*. 2024). Studying young birds is critical for not only optimizing future management strategies but also understanding the consequences of current strategies.

Overall, we recommend that more active monitoring and protection of prebreeders will help support seabird populations. We also highlight three broader themes. First is the importance of long-term population studies for conservation (Lindenmayer *et al*. 2012). Much of the limited information on prebreeders comes from decades-long efforts to band and recapture individual birds (see sources in data file), a difficult and declining practice. Second is the importance of basic behavioral studies. Fundamental ethological questions about seabird behavior (why do young birds delay reproduction?) are in fact tied to urgent questions about conservation (how could we get young birds to start breeding?). Finally, life history characteristics—such as elasticities and breeding ages—should not be viewed as immutable, evolutionary relicts. Instead, life histories are dynamic, ecologically plastic, and critically relevant to protecting species.

## Acknowledgements

We are grateful to Stephen Stearns, Arata Honda, David Wilson, and members of the Cognitive and Behavioural Ecology Program at Memorial University for comments that improved the manuscript. This work is inspired by the fieldworkers who spend decades monitoring seabird populations on remote islands.

