## Appendix S1 for "Prebreeding populations and the importance of life history for conserving the world’s imperiled seabirds"

for

Prebreeding populations and the importance of life history for conserving the world’s imperiled seabirds

L. U. Taylor and E. Gnam

**Table of Contents**

| pp. | 2–5 | Table S1 |
| --- | --- | --- |
| p. | 6 | Table S2 |
| p. | 7 | Figure S1 |
| p. | 8 | Figure S2 |
| pp. | 9–11 | Panel S1 |

**Table S1.** Example studies (n = 31) using remote tracking technologies to monitor young seabirds. Technologies included global position systems (GPS), global location sensors (GLS), satellite transmitters, and accelerometers. We defined age classes as juvenile (J), subadults (S), and breeding adults (A). Juveniles are birds in their first year of life. Subadults are birds between 12 months old and age at first breeding. Sources are arranged chronologically, then alphabetically by first author.

|  |  |  | **Age classes** | | |
| --- | --- | --- | --- | --- | --- |
|  | **Reference** | **Taxon** | **J** | **S** | **A** |
|  | Weimerskirch et al. (2006) | Snowy Albatross (Diomedea exulans) | X |  |  |
|  | Daunt et al. (2007) | European Shag (Gulosus aristotelis) | X |  | X |
|  | Alderman et al. (2010) | White-capped Alabatross (Thalassarche cauta) | X |  |  |
|  | Votier et al. (2011) | Northern Gannet (Morus bassanus) |  | X |  |
|  | Péron and Grémillet (2013) | Scopoli's Shearwater (Calonectris diomedea) | X | X | X |
|  | Riotte-Lambert and Weimerskirch (2013) | Snowy Albatross (Diomedea exulans) | X | X | X |
|  | Bentzen and Powell (2015) | King Eider (Somateria spectabilis) | X | X |  |
|  | Gutowsky et al. (2014) | Black-footed Albatross (Phoebastria nigripes) | X |  | X |
|  | Puetz et al. (2014) | King Penguin (Aptenodytes patagonicus) | X |  |  |
|  | Blanco et al. (2015) | Southern Giant-Petrel (Macronectes giganteus) | X |  | X |
|  | Fayet et al. (2015) | Manx Shearwater (Puffinus puffinus) |  | X | X |
|  | Mendez et al. (2017) | Red-footed Booby (Sula sula) | X |  | X |
|  | Votier et al. (2017) | Northern Gannet (Morus bassanus) |  | X | X |
|  | Grecian et al. (2018) | Northern Gannet (Morus bassanus) |  | X | X |
|  | Afán et al. (2019) | Scopoli's Shearwater (Calonectris diomedea) | X |  | X |
|  | Corbeau et al. (2019) | Great Frigatebird (Fregata minor) | X |  | X |
|  | Labrousse et al. (2019) | Emperor Penguin (Aptenodytes forsteri) | X |  |  |
|  | Pettex et al. (2019) | Northern Gannet (Morus bassanus) |  | X | X |
|  | Ramos et al. (2019) | Cory's Shearwater (Calonectris borealis) | X |  |  |
|  | Campioni et al. (2020) | Cory's Shearwater (Calonectris borealis) |  | X | X |
|  | Collet et al. (2020) | Great Frigatebird (Fregata minor)  Red-footed Booby (Sula sula) | X |  | X |
|  | Mendez et al. (2020) | Red-footed Booby (Sula sula) | X |  | X |
|  | **Table S1 continued.** |  |  |  |  |
|  | |  | **Age classes** | | |
| **Reference** | | **Taxon** | **J** | **S** | **A** |
| Borrmann et al. (2021) | | Lesser Black-backed Gull (Larus fuscus) | X |  |  |
| Enstipp et al. (2021) | | King Penguin (Aptenodytes patagonicus) | X |  | X |
| Pajot et al. (2021) | | Snowy Albatross (Diomedea exulans)  Amsterdam Albatross (Diomedea amsterdamensis) | X |  | X |
| Frankish et al. (2022) | | Gray-headed Albatross (Thalassarche chrysostoma) | X |  |  |
| Gimeno et al. (2023) | | Yellow-legged Gull (Larus michahellis) | X | X |  |
| Delord et al. (2024) | | Amsterdam Albatross (Diomedea amsterdamensis) | X | X | X |
| Morel et al. (2024) | | Lesser Black-backed Gull (Larus fuscus) | X |  | X |
| Navarro et al. (2024) | | Yellow-legged Gull (Larus michahellis) | X | X | X |
| Ponti et al. (2024) | | Audouin's Gull (Ichthyaetus audouinii) | X |  | X |

**Table S2.** Taxonomic groups of seabirds used in the initial search for data (n = 363 species) and final demographic dataset (n = 84 species). Taxonomy follows Clements checklist, October 2023 version (Clements et al. 2023).

|  |  |  |  | **No. species** | |
| --- | --- | --- | --- | --- | --- |
| **Order** | **Family** | **Genus** | **Common** | **Initial search** | **Final dataset** |
| Anseriformes | Anatidae | Polysticta | Sea ducks | 1 | 1 |
|  |  | Somateria |  | 3 | 3 |
|  |  | Histrionicus |  | 1 | 1 |
|  |  | Melanitta |  | 6 | 1 |
|  |  | Clangula |  | 1 | 0 |
|  |  | Bucephela |  | 3 | 1 |
| Charadriiformes | Stercorariidae | Stercorarius | Skuas and jaegers | 7 | 4 |
|  | Alcidae | (All) | Auks, murres, and puffins | 24 | 6 |
|  | Laridae | (All) | Gulls and terns | 100 | 20 |
| Phaethontiformes | Phaethontidae | Phaethon | Tropicbirds | 3 | 2 |
| Sphenisciformes | Spheniscidae | (All) | Penguins | 18 | 7 |
| Procellariiformes | (All) | (All) | Tubenoses | 142 | 26 |
| Suliformes | Fregatidae | Fregata | Frigatebirds | 5 | 1 |
|  | Sulidae | (All) | Boobies and gannets | 10 | 6 |
|  | Phalacrocoracidae | (All) | Cormorants and shags | 39 | 5 |

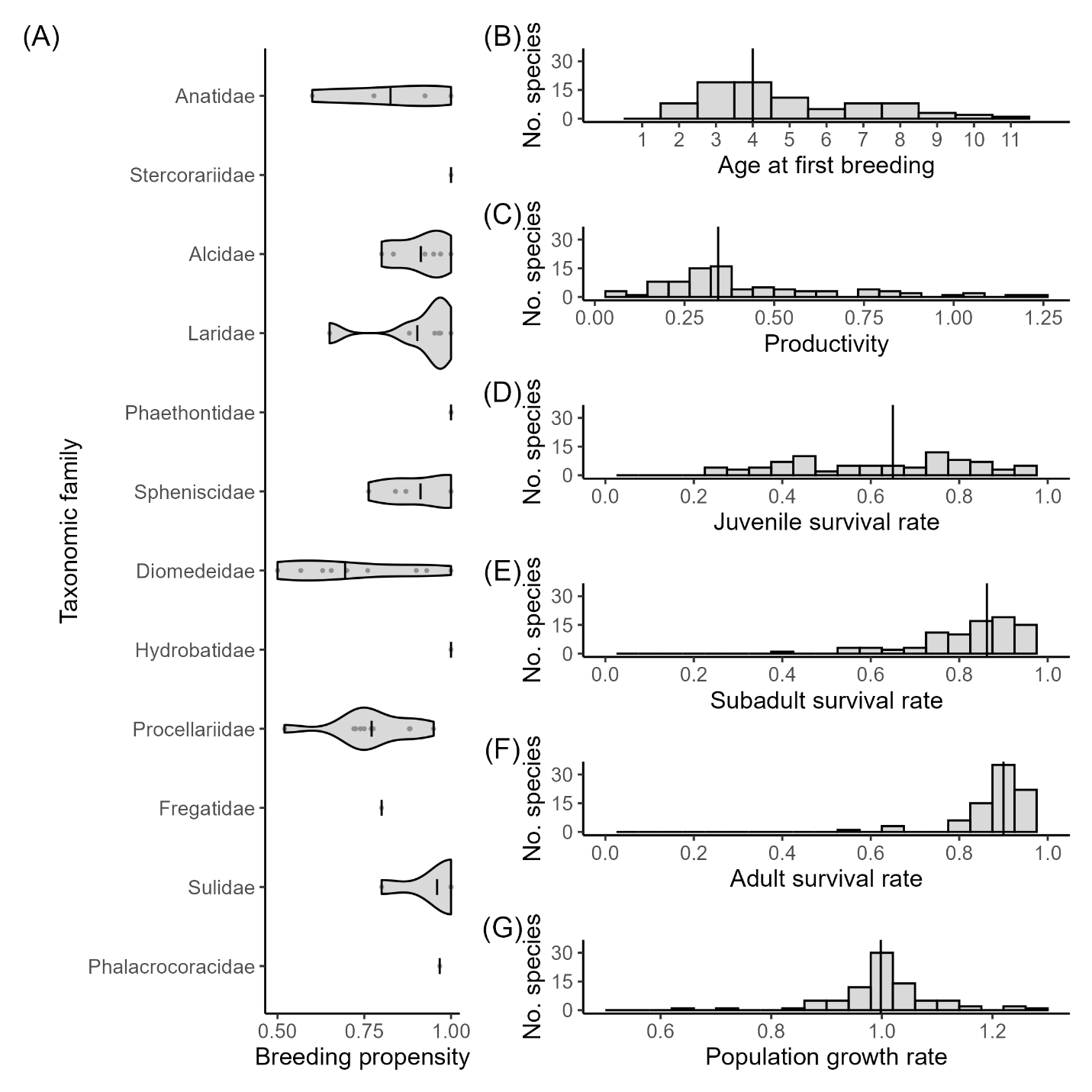

**Figure S1**. Simple demographic parameters across 84 species of seabirds. (A) Annual breeding propensity organized by taxonomic family. Species with unknown breeding propensities were assigned the arithmetic mean value (vertical lines) from species with available data in the same family. (B) Age at first breeding. (C) Productivity. (D) Juvenile survival rate. (E) Subadult survival rate. (F) Adult survival rate. (G) Population growth rate, calculated from the Leslie matrix of each species. Vertical lines in histograms show median.

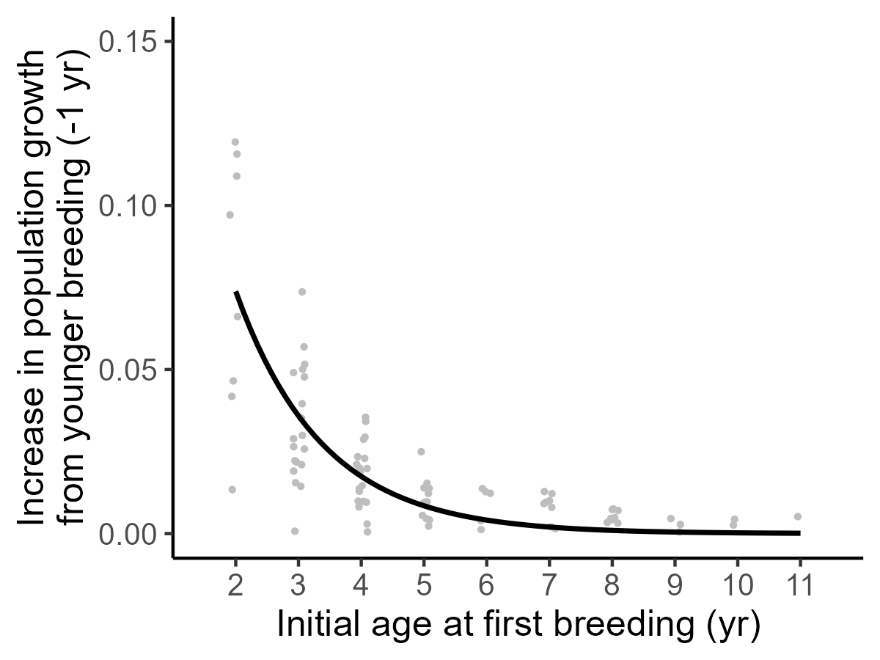

**Figure S2**. Seabirds with lower ages at first breeding show a greater increase to population growth rate given a hypothetical, one-year decrease to age at first breeding. Points slightly jittered along the x-axis. Black curve shows fit from a significant exponential model (a = 0.31, b = -0.72, P < 0.001, AIC = -456), which was preferred to a significant linear model (intercept = 0.06, slope = -0.01, P < 0.001, AIC = -412).

**Panel S1.** Building and analyzing Leslie matrices in this study, including important simplifying assumptions.

For each seabird species in our dataset, we constructed a Leslie matrix to encode population-level breeding and survival rates (Caswell 2001). A Leslie matrix is a simple format that allows one to project population dynamics, such as population growth rate, from underlying demographic parameters. Matrix parameters included fecundity (F), juvenile survival rate (S_J_), subadult survival rate (S_S_), and adult survival rate (S_A_). Fecundity is the product of productivity (P) and breeding propensity (B). Age at first breeding determines the number of rows and columns in the matrix (i.e., the number of stages an individual must pass through, with survival rate S_S_, before becoming a breeding adult). For a species that begins breeding at age two, a Leslie matrix takes the form:

$$\begin{matrix} 0 & 0 & F=P*B \\ S_{j} & 0 & 0 \\ 0 & S_{S} & S_{A} \end{matrix}$$

Population growth rate (λ) is estimated as dominant real eigenvalue of the matrix, and stable age distribution is estimated from the right eigenvector (Caswell 2001). For a parameter in row i and column j of the matrix (A_i,j_), sensitivity quantifies the small change in population growth rate given a small change in that parameter. Sensitivities are calculated from the matrix via products of the right eigenvector (stable age distribution, w) and the left eigenvector (v, reproductive value):

$$Sensitivity_{A_{i,j}}=\frac{d\lambda}{dA_{i,j}}=v_{i}w_{j}$$

Relative sensitivities, called elasticities, are obtained by scaling sensitivity values in terms of parameter magnitude and population growth rate:

$$Elasticity_{A_{i,j}}=\frac{A_{i,j}}{\lambda}*Sensitivity_{A_{i,j}}$$

Because we used simple Leslie matrices—where only the terminal, adult stage had nonzero fecundity—there were only two unique elasticities for each matrix: (1) elasticity for S_A_ and (2) a shared value for all of F, S_J_, and S_S_ (Caswell 2001 pp. 231–2).

To illustrate the impact of potential, future demographic changes, we calculated elasticities before and after changing the underlying parameters. For example, the matrix on the left was used to calculate the original elasticities for Marbled Murrelet (Brachyramphus marmoratus; Skrabis 2005), while the modified matrix on the right was used to calculate hypothetical elasticities after a 20% increase to fecundity:

$\begin{matrix} 0 & 0 & 0.112 \\ 0.62 & 0 & 0 \\ 0 & 0.72 & 0.82 \end{matrix}$ vs. $\begin{matrix} 0 & 0 & 0.112*1.20 \\ 0.62 & 0 & 0 \\ 0 & 0.72 & 0.82 \end{matrix}$

To illustrate the impact of changing the age at first breeding, we modified the structure of each matrix. For example, these two matrices were used to calculate the original population growth rate for Marbled Murrelet (left) along with hypothetical growth rate given a one-year decrease in age at first breeding (right):

$\begin{matrix} 0 & 0 & 0.112 \\ 0.62 & 0 & 0 \\ 0 & 0.72 & 0.82 \end{matrix}$ vs. $\begin{matrix} 0 & 0.112 \\ 0.62 & 0 \\ 0 & 0.82 \end{matrix}$

Importantly, this comparison assumes no consequence, or tradeoff, associated with breeding at a younger age; individuals simply progress more quickly from juveniles to breeding adults. Field studies suggest these tradeoffs exist, with individuals that begin breeding at younger ages also experiencing reduced fecundity or survival (e.g., Pyle et al. 1997; Kim et al. 2011). Because we neglect such tradeoffs, our estimates of the benefits of reduced breeding age on population growth rate are likely biased upward.

Indeed, our simple matrix formulation neglected several important axes of demographic variation. We used broad age categories (i.e., juvenile, subadult, breeding adult), and one characteristic value for each parameter per species. In contrast, field studies show substantial variation across years (e.g., Frederiksen et al. 2008), within age classes (e.g., Weimerskirch 1992), and among populations (e.g., Genovart et al. 2018). Understanding this variation is key to investigating the demographic dynamics of individual species. Nevertheless, the breadth of intraspecific variation is likely encompassed by the broader, interspecific variation spanning multiple seabird orders.
